# β-L-1-[5-(E-2-Bromovinyl)-2-(Hydroxymethyl)-1,3-Dioxolan-4-yl)] Uracil (L-BHDU) Inhibits Varicella Zoster Virus Replication by Depleting the Cellular dTTP Pool

**DOI:** 10.1101/2020.02.13.948216

**Authors:** Chandrav De, Dongmei Liu, Uma S. Singh, Chung K. Chu, Jennifer F. Moffat

## Abstract

ß-L-1-[5-(E-2-bromovinyl)-2-(hydroxymethyl)-1,3-(dioxolan-4-yl)] uracil (L-BHDU) inhibits varicella zoster virus (VZV) replication in cultured cells, human skin, and in SCID-Hu mice with skin xenografts. VZV thymidine kinase converts L-BHDU to monophosphate (MP) and diphosphate (DP) forms, but the triphosphate form was not detected in infected cells and the antiviral mechanism was unknown. Given its similar structure to uridine, we asked if L-BHDU interfered with viral DNA replication via inhibition of the purine and/or pyrimidine biosynthesis pathways. Addition of purines to the medium was unable to restore VZV replication in the presence of L-BHDU. In contrast, excess thymidine and uridine in proportion to L-BHDU restored VZV replication, suggesting that the active form of L-BHDU interfered with pyrimidine biosynthesis. However, addition of thymidine and uridine failed to restore VZV replication in non-dividing cells treated with L-BHDU. Like other herpesviruses, VZV infection increased thymidine triphosphate (dTTP) in confluent cells while L-BHDU treatment decreased the dTTP pool by nearly 4-fold. The active form(s) of L-BHDU did not interfere with cellular metabolism, suggesting viral target(s).

## 1. Introduction

Varicella zoster virus (VZV) is a common α-herpesvirus that causes varicella (chicken pox) upon primary infection. VZV establishes life-long latency in dorsal root and trigeminal ganglia, and reactivation results in herpes zoster (shingles) (Reviewed in Zerboni et al., 2014). Acyclovir (ACV) and its derivatives, foscarnet (phosphonoformate, PFA), and brivudine (BVdU) are antiviral drugs used to treat VZV infections. We found that VZV is highly susceptible to a novel uridine analogue, ß-L-1-[5-(E-2-bromovinyl)-2-(hydroxymethyl)-1,3-(dioxolan-4-yl)] uracil ^(L-BHDU) (De et al., 2014), but its mechanism of action is unknown.^ The antiviral activity of L-BHDU is dependent on viral thymidine kinase (TK), and amino acid substitutions in active domains of the enzyme render the virus resistant to L-BHDU. VZV TK phosphorylates L-BHDU to monophosphate (MP) and diphosphate (DP) forms, but the triphosphate (TP) form was not detected (manuscript in preparation). The general mode of action of antiviral nucleoside analogues is through competitive inhibition of viral DNA polymerase (pol) and/or DNA chain termination (Sauerbrei et al., 2011). Interestingly, the active forms of common antiviral compounds differ. The acyclic nucleoside analogues acyclovir, ganciclovir, penciclovir, famciclovir and brivudine are active in their TP form (De Clercq, 2004; Freeman and Gardiner, 1996). Trifluridine and sorivudine in the MP form inhibit thymidylate synthase (De Clercq, 1996; Hideki Kawai et al., 1993).

Viral replication is tightly linked with the cellular nucleotide metabolism. Herpesviruses encode proteins to facilitate DNA replication when cellular dNTPs are limited, especially deoxythymidine triphosphate (dTTP). There are two pathways for biosynthesis of dTTP in the cell. One pathway is via the production of dTMP from deoxyuridine monophosphate (dUMP), which is anabolized by thymidylate synthase (TS), and the other is via thymidine by thymidine kinase (TK) (Hoffmann et al., 2011; H Kawai et al., 1993). Recent and past studies highlight that components of the pyrimidine biosynthesis pathway represent potential therapeutic targets (Fischer et al., 2013; Hoffmann et al., 2011; McLean et al., 2001). The purpose of this study was to determine the role of L-BHDU in nucleotide metabolism pathways and decipher the possible antiviral mode of action of L-BHDU. We found that L-BHDU interfered with pyrimidine biosynthesis, possibly by targeting viral enzymes, and inhibited VZV DNA replication by depleting the cellular dTTP pool.

## 2. Materials and Methods

### 2.1. Propagation of cells and viruses

Human foreskin fibroblasts (HFFs) (CCD-1137Sk; American Type Culture Collection, Manassas, VA), used prior to passage 20, and human melanoma cell line (MeWo) (ATCC HTB-65) were grown in Eagle minimum essential medium with Earle’s salts and L-glutamine (HyClone Laboratories, Logan, UT), supplemented with 10% heat-inactivated fetal bovine serum (Benchmark FBS; Gemini Bio Products, West Sacramento, CA), penicillin-streptomycin (5000 IU/mL), amphotericin B (250 lU/mL), and nonessential amino acids (all Mediatech, Herndon, VA). ARPE-19 human retinal pigment epithelial cells were grown in Dulbecco’s modified Eagle Medium (DMEM F-12) with 10% fetal bovine serum, penicillin-streptomycin (5000 IU/mL) and amphotericin B (250 lU/mL). Cell were cultured in humidified CO_2_ (5%) at 37°C. VZV-BAC-Luc (Zhang et al., 2007), derived from the Parental Oka (POka, Accession number: AB097933) strain and VZV 13S were propagated in HFFs. VZV 13S was a kind gift from Dr. Jeffery Cohen, NIH (Cohen and Seidel, 1993) that does not express viral thymidylate synthase (ORF 13).

### 2.2. Compounds

L-BHDU was synthesized as described before (Choi et al., 2000). (E)-5-(2-bromovinyl)-2′-deoxyuridine (BVdU, B9647), thymidine (T1895), uridine (U3003), cytidine (C4654), orotate (O2875), dihydroorotate (DHO, D7003), adenosine (A4036), inosine (I4125) and guanosine (G6264) were purchased from Sigma Aldrich, St. Louis, MO. Stock solutions of L-BHDU and BVdU were prepared in dimethyl sulfoxide (DMSO, D2650; Sigma Aldrich), aliquoted and stored at −20°C. The stock solutions of the nucleoside bases were prepared in water, aliquoted and stored at −80°C. Working solutions were prepared fresh for each experiment. dTTP, dCTP and dUTP were purchased from Thermo Scientific (R0181 and R0133). dTMP (T7004), dUMP (D3876) and dTDP (T9375) were purchased from Sigma Aldrich.

### 2.3. VZV rescue experiment in dividing cells

HFFs cells were seeded in clear bottom, black-sided, 6-well plates (W1150, Genetix, Molecular Devices) 24h prior to infection. Dividing cells were infected with cell-associated VZV-BAC-Luc and VZV 13S showing more than 80% cytopathic effect (CPE) at 1:100 ratio of infected to uninfected cells and adsorbed for 2 h at 37°C. Excess virus was removed and the cells were washed once with PBS. Medium containing either vehicle or 2-fold dilutions of the test compounds at concentrations between 0.25 and 2.0 µM were added; this point was deemed time zero. The nucleoside supplements were added at a constant concentration of 200 µM together with different concentrations of L-BHDU. Cells were treated for 48 h and the medium containing the drug and/or nucleoside supplements was changed after 24 h. VZV-BAC-Luc yield was determined by bioluminescence imaging using the IVIS^®^ 50 instrument (Caliper Life Sciences/Xenogen, Hopkinton, MA) and expressed as Total Flux (photons/sec/cm^2^/steradian) at 48 hpi (De et al., 2014). Virus yield for VZV 13S and VZV-BAC-Luc was also measured using a quantitative real time PCR method described previously (De et al., 2014).

### 2.4. VZV rescue experiment in non-dividing cells

HFFs were seeded in clear bottom, black-sided, 6-well plates (W1150, Genetix, Molecular Devices). Cells were grown in 10% FBS medium for three days until they were contact inhibited. After 72 h, the serum in the medium was reduced to 5% for an additional 48 h. After five days of cell culture, the non-dividing HFFs were infected with cell-associated VZV-BAC-Luc and VZV_13S_ at 1:100 ratio of infected to uninfected cells. The rescue experiment was performed in the same manner as in 2.3.

### 2.5. MTT assay for HFF-TK cell proliferation

HFF-TK cells were generated as described previously (manuscript in preparation) and grown in the same manner as HFF cells with the addition of puromycin (2 µg/mL) to the medium to maintain selection for the integrated lentivirus expression vector. HFF-TK cellular proliferation was evaluated by colorimetric MTT (3-(4,5-dimethylthiazol-2-yl)-2,5-diphenyl tetrazolium bromide) assay (Mosmann, 1983). In brief, the cells were seeded 1:3 in 48-well plates and incubated at 37°C in 5% CO_2_ overnight. The cells were treated with antiviral drugs and/or pyrimidine bases in eight replicates.

### 2.6. Measurement of dNTPs by LC-MS/MS

ARPE-19 cells were grown in 175-cm^2^ flasks and harvested at different time points and conditions. Dividing ARPE-19 cells were harvested 27 h after plating. Non-dividing ARPE-19 cells were harvested 3 days after confluence was reached. Duplicate flasks of confluent ARPE-19 cells were infected with cell-associated VZV-BAC-Luc at 1:20 ratio of infected to uninfected cells. One infected culture of ARPE-19 cells was treated with L-BHDU (2 µM) after 24 h. Both infected cultures of ARPE-19 cells were harvested after 48 h. The ARPE-19 cell monolayer was scraped, pelleted, and washed three times with PBS. The cells were suspended in chilled acetonitrile (50% in water) for 10 min on ice. The cell suspensions were centrifuged at 18,000x*g* at 0°C for 5 min. The supernatants were collected and dried in a lyophilizer. The dried extracts were suspended in deionized water and stored at −80°C. The cell lysates were then processed for dNTP detection by LC-MS/MS. Selected reaction monitoring (SRM) was used to detect and quantify the following deoxynucleotide phosphates: dTMP, dTDP, dTTP, dUMP, dUTP and dCTP. SRM transitions were determined by teeing 20 µL of dNTP standard (100 µM in water with 380 µL of 50:50 Solvent A: Solvent B, such that the final concentration of the standard was 5 µM when infused into the instrument. LC-MS analysis was performed using a microspray triple quadrupole system. Samples were separated on a porous graphite column (Hypercarb 100x 2.1 mm column, ThermoScientific) using a Dionex Ultimate 3000 HPLC coupled to a Quantum Access Max mass spectrometer (Thermo Scientific) and run in the SRM negative scan mode. The solvent system was 2 mM ammonium acetate (pH 10, Solution A) and 100% acetonitrile (Solution B) in a 6.5 minute HPLC method. Using a flow rate of 400 µL/min, the column was equilibrated at 40°C in 2% Solution B for 1 min, ramped to 25% Solution B in 2.5 min, ramped to 70% Solution B in 1 min, and then held there for 1 min before being returned to the equilibration condition of 2% Solution B for the remainder (2 min) of the run. Analysis of the extracted ion chromatograms and peaks was performed with the Thermo Scientific LC Quan and XCalibur software. 5 point smoothing using the Boxcar method and integration using Thermo Scientific’s “ICIS” algorithm were employed.

### 2.7. Statistical analysis

GraphPad Prism 5.02 for Windows (Graph-Pad Software, San Diego, CA, www.graphpad.com) was used for statistical calculations. A *p* ≤ 0.05 was considered statistically significant.

## 3. Results

### 3.1. Pyrimidines rescued VZV from L-BHDU in primary cells

Nucleoside analogues often affect nucleotide metabolism (Galmarini et al., 2001; Parker, 2009), and so we hypothesized that the active form of L-BHDU might prevent VZV replication by this mechanism. To address this question, VZV cultures in two different cell types were treated with L-BHDU and various purine and pyrimidine bases were added in excess to determine whether they could rescue virus replication (Fig.1.A). If a purine or pyrimidine rescues virus replication, then that pathway is likely blocked by the active form of the drug. In this assay, the amount of purine or pyrimidine was held constant while the drug concentration was serially decreased 2-fold. Purines did not rescue viral replication in the presence of L-BHDU. Conversely, addition of the pyrimidines thymidine and uridine rescued VZV replication and reversed the inhibitory effect of L-BHDU in a dose dependent manner. A 200:1 ratio of thymidine to L-BHDU restored VZV replication while uridine required 800:1 for the same effect. This was observed in HFFs, primary dermal fibroblasts, (Fig.1.A) but not in MeWos, a melanoma cell line (Fig.1.B). Dihydroorotate (DHO) and orotate (intermediates in the pyrimidine *de novo* synthesis pathway) did not rescue VZV replication. This suggested that L-BHDU blocked the pyrimidine salvage pathway in VZV-infected cells.

**Figure 1.**
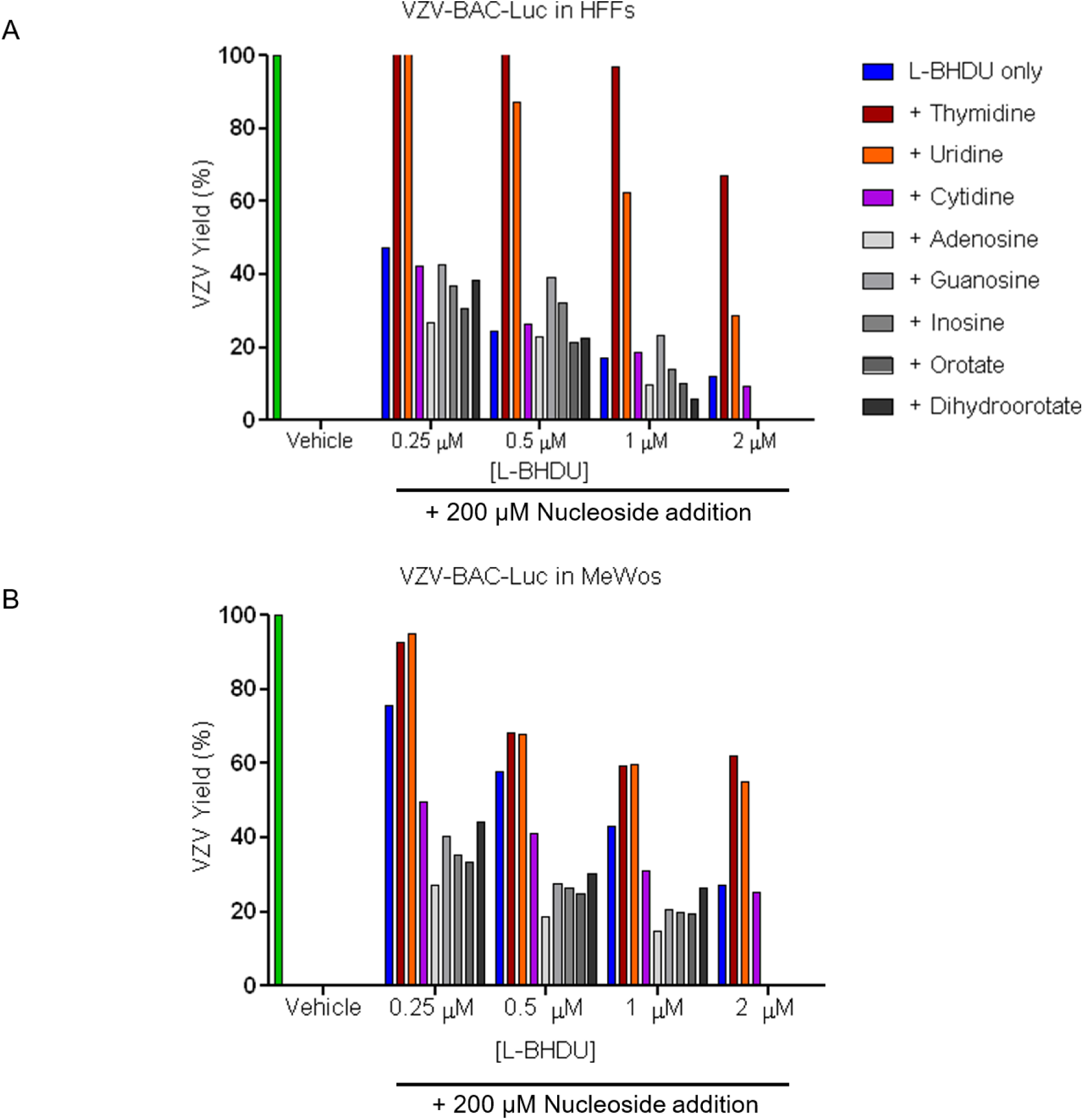
Pyrimidines thymidine and uridine rescued VZV from inhibition by L-BHDU. Dividing HFFs (A) or MeWo cells (B) were infected with VZV and treated with L-BHDU from 0.25 to 2.0 µM (approximately EC_50_ to EC_99_ concentrations) and various purine and pyrimidine pathway intermediates were added to the medium at 200 µM. VZV yield was measured by bioluminescence imaging and normalized to the vehicle group (set at 100%) for each cell type. Data are representative of at least three separate experiments done in triplicates.

### 3.2. Cell proliferation enzymes and VZV ORF13 (thymidylate synthase) were required for pyrimidines to rescue VZV from L-BHDU

The abundance of nucleotides in a cell, known as the nucleotide pool, varies greatly during the cell cycle. Nucleotides are scarce in confluent or quiescent cells whereas dividing cells require high concentrations of dNTPs for genome synthesis during S phase (Hollenbaugh et al., 2013). Herpesviruses use dNTPs for viral DNA replication, thus they encode metabolic enzymes involved in nucleotide metabolism (Vastag et al., 2011). It was not known whether the antiviral effects of L-BHDU could be overcome by exogenous pyrimidines in confluent cells, in which the enzymes required for dTTP synthesis are not expressed. VZV grew normally in confluent HFFs and was inhibited in the presence of L-BHDU (Fig. 2.B and D). This antiviral effect was not rescued by the addition of excess thymidine or uridine, in contrast to the results in dividing cells. VZV is one of two human herpesviruses other than HHV8 that encodes a viral thymidylate synthase (TS) (Gáspár et al., 2002), thus cellular TS may not be required in confluent cells.

**Figure 2.**
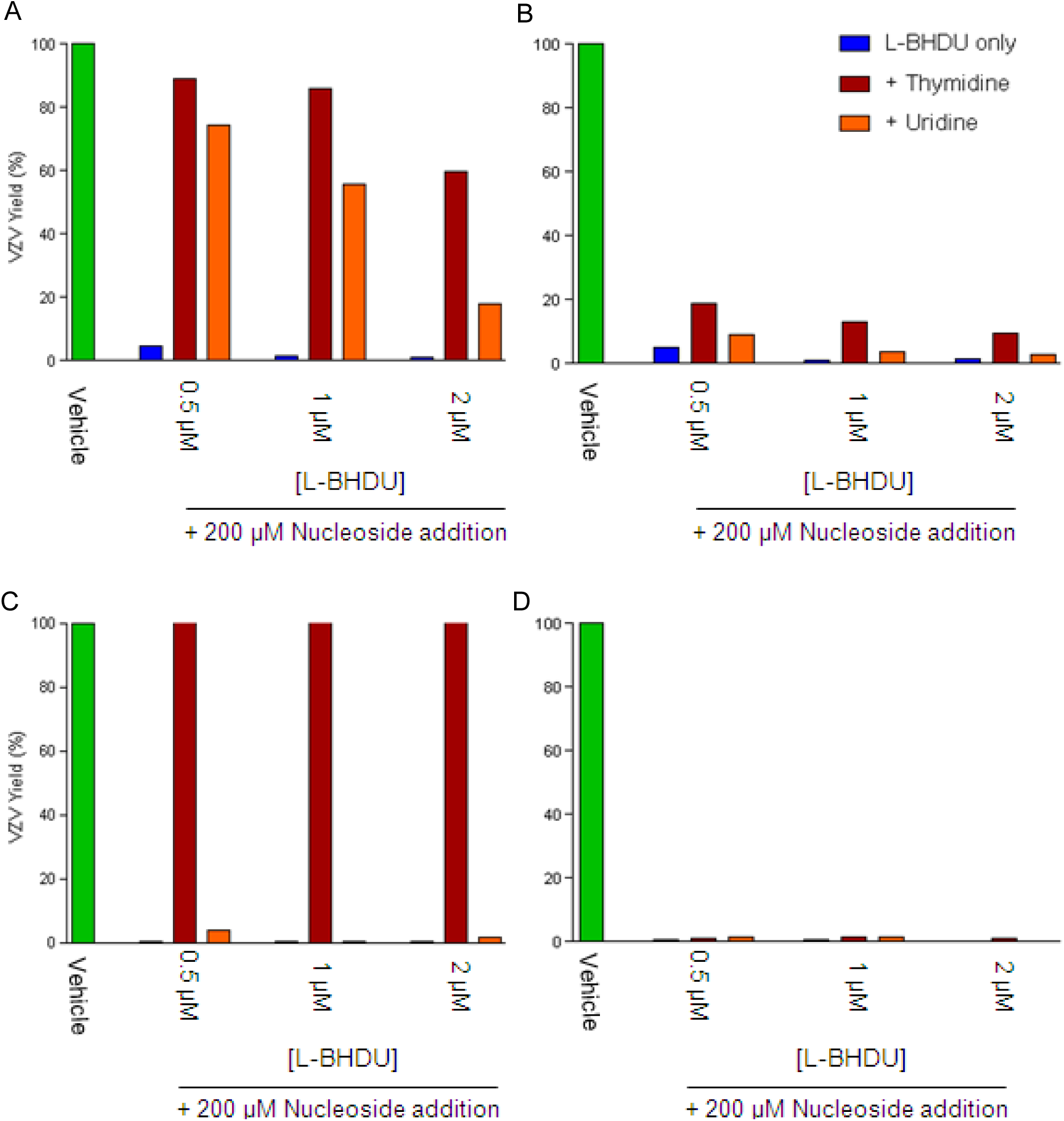
Thymidine and uridine failed to rescue VZV replication from L-BHDU inhibition in non-dividing cells. Dividing (A, C) or confluent (B, D) HFFs were infected with VZV-BAC-Luc (A, B) or VZV 13S (C, D) and treated with L-BHDU from 0.5-2.0 µM and thymidine or uridine was added to the medium at 200 µM. VZV yield was measured by bioluminescence imaging (A, B) or qPCR (C, D) and normalized to the vehicle group (set to 100%). Data are representative of at least two separate experiments done in triplicates.

Indeed, the TS gene encoded by ORF13 is not essential and a stop codon mutant, VZV13S, grows normally (Cohen and Seidel, 1993). TS is essential for regulating the balanced supply of dNTP precursors for DNA replication (Kaneda et al., 1990). Therefore, the intracellular availability of dUMP may be the rate-limiting step in dTMP biosynthesis and hence critical for efficient viral replication in quiescent cells (Gribaudo et al., 2003). We tested whether addition of thymidine or uridine could rescue VZV 13S replication from L-BHDU inhibition in dividing and non-dividing cells. As expected, VZV 13S grew well in both dividing and non-dividing cells in the absence of L-BHDU. In non-dividing cells neither thymidine nor uridine restored VZV 13S replication (Fig. 2.D). Interestingly, thymidine but not uridine restored VZV 13S replication in dividing cells (Fig. 2.C). Thus, viral TS was important for utilizing uridine to overcome the effects of L-BHDU. Furthermore, cellular enzymes in proliferating cells were also necessary to rescue WT and 13S VZV from L-BHDU.

### 3.3. Active L-BHDU did not inhibit cell proliferation

To determine whether the active form of L-BHDU affected cellular metabolism, a proliferation assay was performed. L-BHDU is phosphorylated to its active form by viral thymidine kinase (TK) and not by cellular TK, thus we used a recombinant HFF-TK cell line stably expressing VZV TK to produce L-BHDU-MP and -DP. BVdU was a positive control because its monophosphorylated form is cytostatic in cells expressing viral TK. BVdU-MP inhibits cellular thymidylate synthase and subsequently cellular proliferation (Balzarini et al., 1994, 1985). HFF-TK cells grew normally in medium and with vehicle (Fig.3). As expected, serum starvation and BVdU treatment significantly inhibited cell proliferation after 72 h. Addition of thymidine but not uridine restored proliferation in the cells treated with BVdU (Fig.3), which reflects the known blockade of the *de novo* biosynthesis pathway by BVdU-MP inhibition of cellular TS. Conversely, HFF-TK cells grew normally in the presence of L-BHDU at the EC_90_ concentration, suggesting that L-BHDU-MP and/or -DP did not inhibit cellular nucleotide metabolism.

**Figure 3.**
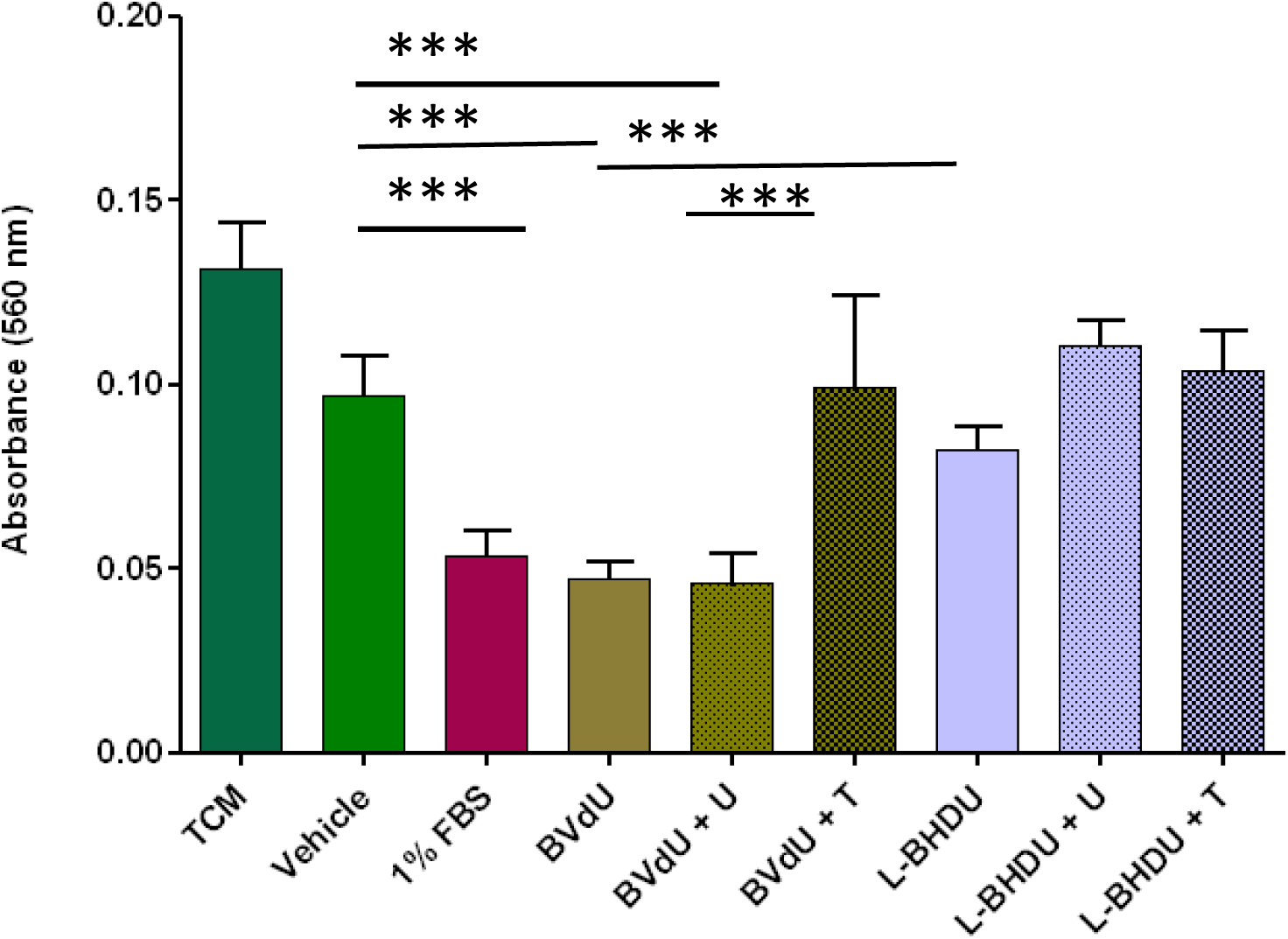
Active L-BHDU did not inhibit HFF-TK proliferation. HFF-TK cells were plated at low density and grown for 72 h with BVdU or L-BHDU. Uridine (U) or thymidine (T) was added to some cultures at 200 µM. Cellular proliferation was measured by MTT assay. Each bar represents the mean ± standard deviation of the absorbance values at 560 nm. Data are representative of at least two independent experiments performed in replicates of six. Significant reductions in cellular proliferation compared to the vehicle group are indicated (* *p*<0.05,one-way ANOVA with Bonferroni multiple comparison correction test).

### 3.4. L-BHDU suppressed the increase in dTTP in VZV-infected cells

One response to herpesvirus infection is a marked increase in the level of dTTP in the cell, which is consistent with the need to replicate viral DNA (Daikoku et al., 1991; Vastag et al., 2011). We hypothesized that L-BHDU indirectly inhibits VZV replication by decreasing the cellular dTTP pool, which prevents viral DNA synthesis. The relative amounts of various nucleotides were measured by HPLC-MS/MS in dividing or confluent cells that were infected with VZV and treated with L-BHDU. ARPE-19 retinal pigment epithelial cells were used for this experiment because they are highly permissive for VZV infection even when contact-inhibited, and they have more abundant dNTPs than HFFs (Sloutskin et al., 2013). As expected, nucleotides increased in ARPE-19 cells when they were released from contact inhibition for 1 day (Fig.4, Table 1). When confluent ARPE-19 cells were infected with VZV for 1 day, all the nucleotides increased to varying degrees. The level of dTMP was 4-fold higher than dividing ARPE-19 cells while dTDP and dTTP were similar. The levels of dUMP, dUTP, and dCTP were less than dividing ARPE-19 cells but more than confluent cells. At this point duplicate cultures of VZV-infected ARPE-19 cells were either treated with L-BHDU or the medium was refreshed. In untreated, VZV-infected ARPE-19 cells, the levels of dTDP and dTTP continued to increase while L-BHDU treatment prevented this. The level of dTMP remained high in VZV-infected cells treated with L-BHDU. Thus L-BHDU prevents the accumulation of dTTP in VZV-infected cells and thereby blocks viral DNA replication.

**Figure 4.**
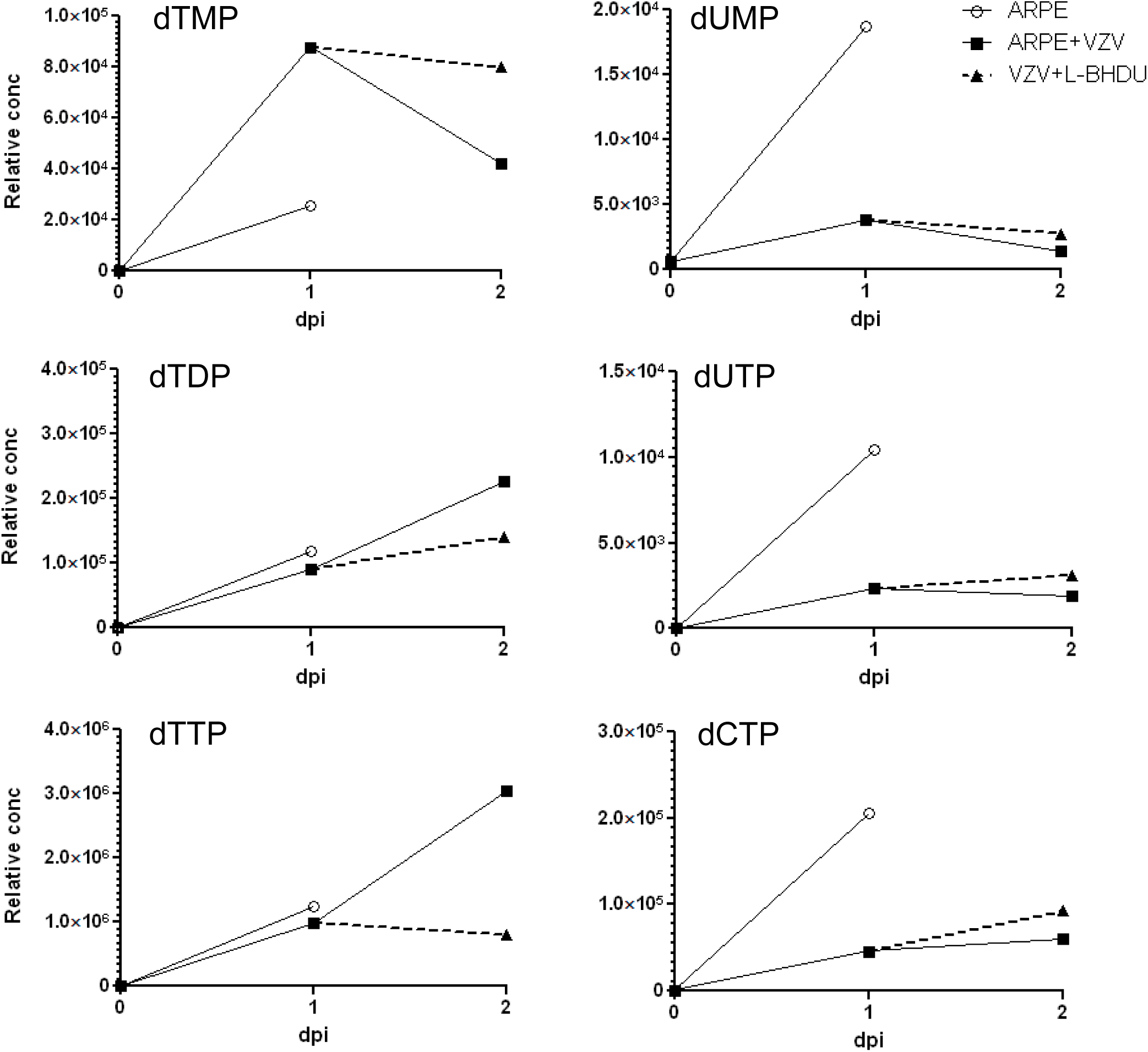
L-BHDU suppressed the increase in dTTP in VZV-infected ARPE-19 cells. Relative concentrations of deoxynucleotide phosphates were measured by LC-MS/MS in confluent and dividing ARPE-19 cells (open circles), VZV-infected confluent ARPE-19 cells (black squares), and VZV-infected, confluent ARPE-19 cells treated with L-BHDU for 24 h (black triangles). Area under the curve (AUC) analysis was used to quantify the relative concentration of the different compounds from the standard curve spanning concentrations of 10 nM to 3.16 µM. Data are representative of two separate experiments performed in triplicate.

**Table 1:** Relative concentrations of deoxynucleotide phosphates in ARPE-19 cells during VZV infection and L-BHDU treatment

| Nucleotide | ARPE-19 cells |  | VZV-infected ARPE-19 cells |  |  |
| --- | --- | --- | --- | --- | --- |
|  | Confluent | Subconfluent | 24 hpi | 48 hpi | L-BH DU 24-48 hpi |
| dTMP | ND* | $2.55 \times 10^4$ | $8.78 \times 10^4$ | $4.21 \times 10^4$ | $7.99 \times 10^4$ |
| dTDP | ND | $1.19 \times 10^5$ | $8.99 \times 10^4$ | $2.26 \times 10^5$ | $1.41 \times 10^5$ |
| dTTP | ND | $1.24 \times 10^6$ | $9.88 \times 10^5$ | $3.04 \times 10^6$ | $7.95 \times 10^5$ |
| dUMP | 630 | $1.87 \times 10^4$ | $3.84 \times 10^3$ | $1.4 \times 10^3$ | $2.76 \times 10^3$ |
| dUTP | ND | $1.04 \times 10^4$ | $2.30 \times 10^3$ | $1.88 \times 10^3$ | $3.10 \times 10^3$ |
| dCTP | 992 | $2.06 \times 10^5$ | $4.58 \times 10^4$ | $5.97 \times 10^4$ | $9.25 \times 10^4$ |

\*ND= Below the level of detection

## 4. Discussion

This study demonstrates that L-BHDU inhibits VZV replication by decreasing the cellular dTTP pool by interfering with the pyrimidine biosynthesis pathway (Fig. 5). There are two routes to the synthesis of pyrimidines; the *de novo* and salvage pathways. Some compounds targeting the *de novo* pyrimidine biosynthesis pathway have antiviral properties (Fischer et al., 2013; Hoffmann et al., 2011; Wang et al., 2011). Our results point away from the *de novo* pathway because addition of intermediates such as orotate and DHO failed to restore VZV replication in the presence of L-BHDU. On the other hand, pyrimidine salvage pathway substrates thymidine and uridine rescued VZV replication from L-BHDU inhibition. The activity of the pyrimidine *de novo* and salvage pathways varies with the developmental stage of the cell. The *de novo* pathway is low in fully differentiated cells, whereas it is indispensable in proliferating cells (Evans and Guy, 2004). Rapidly dividing cancer cells have much higher levels of cellular dNTPs than primary cells (Amie et al., 2013) and this can reduce phosphorylation of antiviral nucleoside analogues (Karlsson et al., 1986). This may explain why L-BHDU was more effective in primary HFF than in MeWo cells (melanoma cell line), and why addition of thymidine and uridine did little to rescue VZV replication in MeWo cells. Addition of thymidine or uridine did not restore VZV replication in confluent HFFs. In non-dividing cells where cellular enzymes are lacking, dNTPs were insufficient for virus replication to resume (Cheng and Traut, 1987; Staub and Eriksson, 2006).

**Figure 5.**
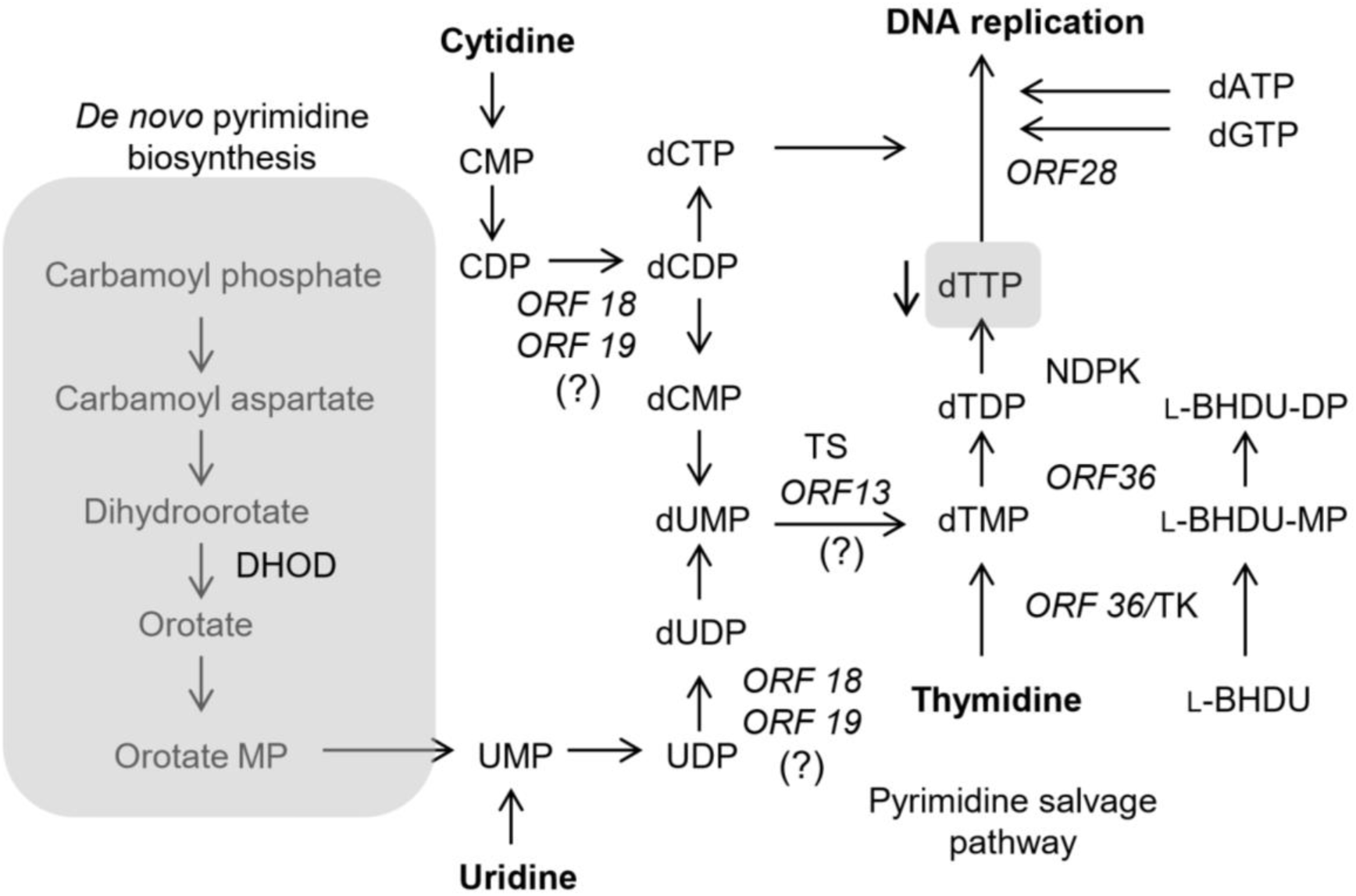
Diagram of the *de novo* and salvage pathways for dNTP synthesis. The de novo pathway and exogenous uridine converge at UMP, which is converted to dTMP by ribonucleotide reductase (RNR) and thymidylate synthase (TS), which each have viral and cellular forms. The pyrimidine salvage pathway uses thymidine to generate the dTTP pool through thymidine kinase (TK, viral and cellular) and nucleotide diphosphate kinase (NDPK, cellular). L-BHDU is phosphorylated by VZV thymidine kinase to mono- and diphosphate forms. It possible that L-BHDU-MP and/or –DP interfere with TS and RNR (noted by ? marks), or by substrate inhibition of viral TK.

Though herpesviruses rely primarily on the metabolic capabilities of their cellular hosts for replication, all herpesviruses encode enzymes that are involved in nucleotide metabolism (Vastag et al., 2011). VZV encodes several enzymes that play critical roles in viral replication, especially those involved in the pyrimidine biosynthesis pathway (Reviewed in Cohen, 2010, 1996). Herpesviruses also actively redirect host cell metabolism to increase the dNTP pool, especially dTTP, by inducing cellular enzymes (Gribaudo et al., 2003, 2002; Vastag et al., 2011). We found that VZV infection of non-dividing cells increased the cellular dTTP pool along with other dNTPs to facilitate viral replication (Fig. 4). Due to the expression of viral enzymes involved in pyrimidine biosynthesis pathway including vTS and increased dNTP pools, VZV replicated well in non-dividing cells.

The maintenance of balanced dNTP pools is critical for viral DNA replication and perturbation can lead to inhibition of virus replication. ACV and bucyclovir (BCV) treatment of HSV-1 infection increase cellular dNTP pools including dTTP (Furman et al., 1982; Karlsson et al., 1986; Prichard et al., 1993). ACV-TP and BCV-TP both target viral DNA polymerase and cause chain termination. This results in reduced utilization of dNTPs and leads to accumulation in treated, infected cells (Furman et al., 1982). Interestingly, L-BHDU lowered the dTTP pool in VZV-infected cells. There several possible explanations for this observation. First, if L-BHDU active form(s) inhibits enzymes in the pyrimidine biosynthesis pathway, then the dTTP pool would be depleted and viral DNA replication would eventually cease. Second, L-BHDU competes with thymidine and dTMP as a substrate of VZV TK. Unlike ACV, L-BHDU might be less sensitive to competition from intracellular thymidine. Third, a higher rate of formation of L-BHDU metabolites compared to the normal substrates (dTMP and dTDP) could lead to a higher accumulation of L-BHDU active forms in the infected cells. These factors could deplete the dTTP pool and prevent virus replication. Depletion of dTTP perturbs the levels of the other deoxynucleotides through feedback mechanisms, and these imbalances severely disrupt cellular DNA synthesis and repair (Longley et al., 2003).

Rescue of VZV replication in the presence of L-BHDU by addition of excess uridine suggests a major role for TS in supplying dTMP via dUMP to restore the cellular dTTP pool. Due to structural similarities between dUMP and L-BHDU-MP, it is possible that L-BHDU-MP might compete with dUMP for the active sites on TS, either viral or cellular. Both cellular and viral thymidylate synthases (cTS and vTS) could be targets of active L-BHDU in infected cells. We found that HFF-TK cells grew normally when active L-BHDU was present, which excludes cTS and other proliferation enzymes as targets. This conclusion was strengthened by our finding that uridine rescued wt VZV in subconfluent HFFs with L-BHDU, but not VZV 13S strain that does not express vTS. This points to the key role of vTS in conversion of dUMP to dTMP to replenish the dTTP pool and suggests that vTS could be a potential target of L-BHDU-MP. However, we found that uridine could not rescue wt or 13S VZV in confluent HFFs in the presence of L-BHDU. The reason why addition of uridine failed to restore VZV replication in confluent cells is due to low translation and activity of uridine kinase and UMP kinase (Cheng and Traut, 1987; Staub and Eriksson, 2006). Without these cellular enzymes, uridine fails to convert to dUMP and vTS lacks its substrate for dTMP synthesis. However, synthesis of dTMP was not sufficient to overcome the antiviral effects of active L-BHDU in confluent cells. Thus there are other targets of active L-BHDU that are necessary for VZV replication in quiescent cells. Ribonucleotide reductase (RNR) catalyzes formation of high concentrations of deoxynucleotides for viral DNA synthesis (Spector et al., 1989, 1987). RNR is essential for virus replication in non-dividing cells (Duan et al., 1998; Sienaert et al., 2002). The natural substrates of RNR are ribonucleoside diphosphates, so it would be interesting to test whether L-BHDU-DP inhibits VZV-encoded RNR. The exact molecular mechanisms that mediate the depletion of dTTP pool by L-BHDU active form(s) have not been fully elucidated and will require further investigation. A complete understanding of the mechanism of action of L-BHDU at the molecular level is an important scientific objective for design and development of more effective antiviral therapy.

## 5. Acknowledgments

Kevin Wallace and Jennifer Hryhorenko at the University of Rochester Proteomics Center performed LC-MS/MS assays to measure cellular dNTP pools. J.F.M. is supported in part by the contract HHSN272201000023I from the Division of Microbiology and Infectious Diseases, NIAID.

